# SARS-CoV-2 infection in free-ranging white-tailed deer (*Odocoileus virginianus*)

**DOI:** 10.1101/2021.11.04.467308

**Authors:** Vanessa L. Hale, Patricia M. Dennis, Dillon S. McBride, Jaqueline M. Nolting, Christopher Madden, Devra Huey, Margot Ehrlich, Jennifer Grieser, Jenessa Winston, Dusty Lombardi, Stormy Gibson, Linda Saif, Mary L. Killian, Kristina Lantz, Rachel Tell, Mia Torchetti, Suelee Robbe-Austerman, Martha I. Nelson, Seth A. Faith, Andrew S. Bowman

## Abstract

Human-to-animal spillover of SARS-CoV-2 virus has occurred in a wide range of animals, but thus far, the establishment of a new natural animal reservoir has not been detected. Here, we detected SARS-CoV-2 virus using rRT-PCR in 129 out of 360 (35.8%) free-ranging white-tailed deer (*Odocoileus virginianus*) from northeast Ohio (USA) sampled between January-March 2021. Deer in 6 locations were infected with at least 3 lineages of SARS-CoV-2 (B.1.2, B.1.596, B.1.582). The B.1.2 viruses, dominant in Ohio at the time, spilled over multiple times into deer populations in different locations. Deer-to-deer transmission may have occurred in three locations. The establishment of a natural reservoir of SARS-CoV-2 in white-tailed deer could facilitate divergent evolutionary trajectories and future spillback to humans, further complicating long-term COVID-19 control strategies.

**One-Sentence Summary:** A significant proportion of SARS-CoV-2 infection in free-ranging US white-tailed deer reveals a potential new reservoir.

## Main Text

To date, SARS-CoV-2, the virus responsible for coronavirus disease 2019 (COVID-19), has caused over 4 million deaths globally (*1*). The zoonotic origins of SARS-CoV-2 are not fully resolved (*2*), leading to active surveillance for susceptible host species and potential new reservoirs. Natural infections of SARS-CoV-2 linked to human exposure have been reported in both domestic animals (e.g. cats, dogs, ferrets) and in wildlife under human care (e.g. several species of big cats, Asian small-clawed otters, western lowland gorillas, and mink) (*3*). Experimental infections have identified additional animal species susceptible to SARS-CoV-2 including hamsters, North American raccoons, striped skunks, white-tailed deer, raccoon dogs, fruit bats, deer mice, domestic European rabbits, bushy-tailed woodrats, tree shrews, and multiple non-human primate species (*4–13*). Moreover, several naturally or experimentally infected species were capable of intra-species SARS-CoV-2 transmission (cats, ferrets, fruit bats, hamsters, raccoon dogs, deer mice, white-tailed deer) (*6–8, 10, 14–16*). Vertical transmission has also been suggested to occur in experimentally infected white-tailed deer (*16*). An *in silico* study modeling ACE2 receptor binding residues across host species predicted cetaceans, rodents, primates, and cervids are at highest risk for infection due to predicted binding between the ACE2 receptor and the SARS-CoV-2 spike protein (*17*).

Limited evidence for SARS-CoV-2 infection or exposure (antibodies) in free-ranging animal populations has also begun to emerge. In Utah (USA), free-ranging mink presumed to be escapees from a mink farm tested positive for SARS-CoV-2 antibodies, and a subset tested positive via rRT-PCR for SARS-CoV-2 (*18*). In Spain, two free-ranging mink also tested positive for SARS-CoV-2 (*19*). Additionally, antibodies for SARS-CoV-2 have been identified in 152 free-ranging white-tailed deer (seroprevalence 40%) sampled across Michigan, Pennsylvania, Illinois, and New York (USA) (*20*). The presence of SARS-CoV-2 in free-ranging animals raises critical concerns about the potential establishment of a new reservoir, future spillback to humans and spillover to other species including livestock based on prior studies of experimental transmission of other betacoronaviruses from cervids to cattle (*21*).

White-tailed deer are of particular interest because their ACE2 receptor binding residues are predicted to have high affinity for SARS-CoV-2 (*17*); they demonstrate experimental susceptibility to and transmission of SARS-CoV-2 (*7, 16*); and, SARS-CoV-2 antibodies have been reported in free-ranging deer (*20*). In this study, we report the presence of SARS-CoV-2 in 129 out of 360 (35.8%) free-ranging white tailed deer (*Odocoileus virginianus*) from northeast Ohio tested via rRT-PCR between January-March 2021. Genetic sequence data was used to estimate the number of human-to-deer transmission events, characterize the genetic diversity of the virus in deer, and identify phylogenetic clades of deer-only viruses consistent with deer-to-deer transmission in certain locations.

## Results

We sampled 360 free-ranging white-tailed deer across from nine locations (Figure 1) in northeast Ohio (USA) between January-March 2021. Across the entire study, SARS-CoV-2 was detected by rRT-PCR in 35.8% of nasal swabs from white-tailed deer (129/360, 95% CI 30.9%, 41.0%, Table S1). Prevalence estimates varied from 13.5% to 70% across the nine sites (Figure 1). Each site was sampled 1-3 times during the study period, for a total of 18 collection dates (Table S2).

**Figure 1.**
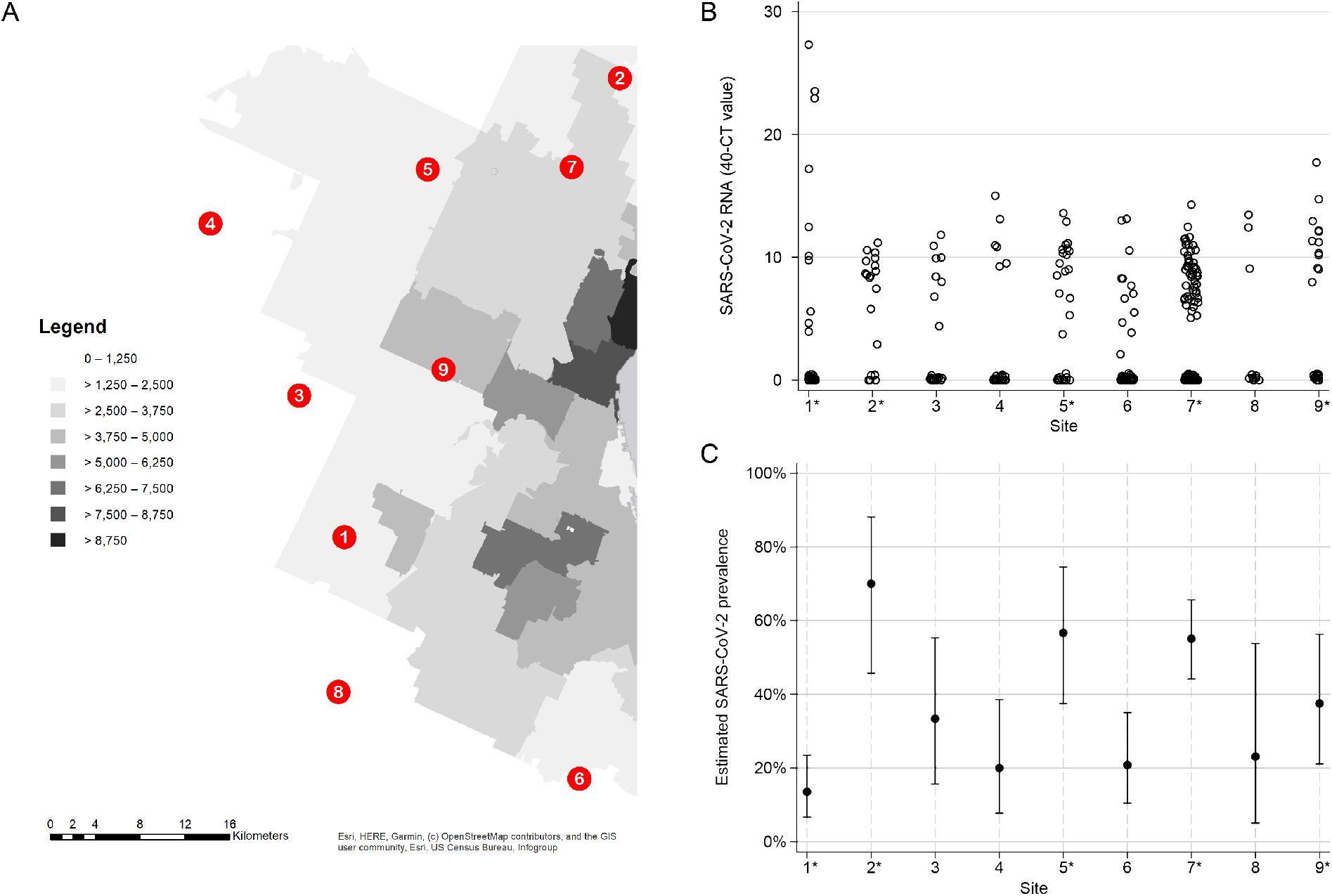
SARS-CoV-2 viral RNA in white-tailed deer across the study locations. (A) The nine study sites were spread across a 1000 km^2^ landscape of varying population density in Northeast Ohio. Darker shading corresponds to higher human population density (people per square mile). Sampling sites one, two, five, seven, and nine are in close proximity to human populations and are indicated as urban sites with an asterisk in panels B and C. (B) Nasal swabs from white-tailed deer were tested for the presence of SARS-CoV-2 viral RNA using real-time reverse transcriptase PCR (rRT-PCR). Estimates of SARS-CoV-2 viral RNA are represented by the Ct value of the N1 rRT-PCR target subtracted from 40. Negative samples are represented with a value of zero. (C) The prevalence of SARS-CoV-2 in the white-tailed deer at each study site was estimated using rRT-PCR (Clopper-Pearson exact 95% confidence interval bars).

At least 1 rRT-PCR-positive sample was identified from 17/18 collection dates. Fewer male deer (n=149) were sampled than female deer (n=211), but males (Chi2 = 25.45, p-value < 0.0005) and heavier deer (Wilcoxon-Mann-Whitney p-value = 0.0056) were more likely to test positive for SARS-CoV-2 (Table S3). Other potential predictors including deer age and harvest method (bait site versus opportunistically) were not associated with SARS-CoV-2 detection (Table S3). The highest prevalence estimates of SARS-CoV-2 were observed in four sites that are more urban and surrounded by higher human population densities (Figure 1).

We obtained 14 whole-genome sequences from six of the nine sites representing seven different collection dates (from 1/26/2021 to 2/25/2021, Table S1). The genetic composition of the deer sequences reflected the dominant lineages that were circulating in humans in Ohio during January-February 2021 (Figure 2, Table S4). The dominant B.1.2 viruses were identified in deer at sites 4, 7, 8, and 9. B.1.596 sequences were identified at site 1 at both collection times. A rarer lineage, B.1.582, was identified at site 6. No sequences consistent with Alpha (B.1.1.7) or Delta (B.1.617.2) were identified in the deer samples; these variants became widespread among the human population only after February 2021. The deer samples were collected several weeks after the peak of the winter epidemic of SARS-CoV-2 in humans in Ohio, when the probability of human-to-deer transmission would have been highest, based solely on the amount of virus in circulation in humans and potentially the environment at that time.

**Figure 2.**
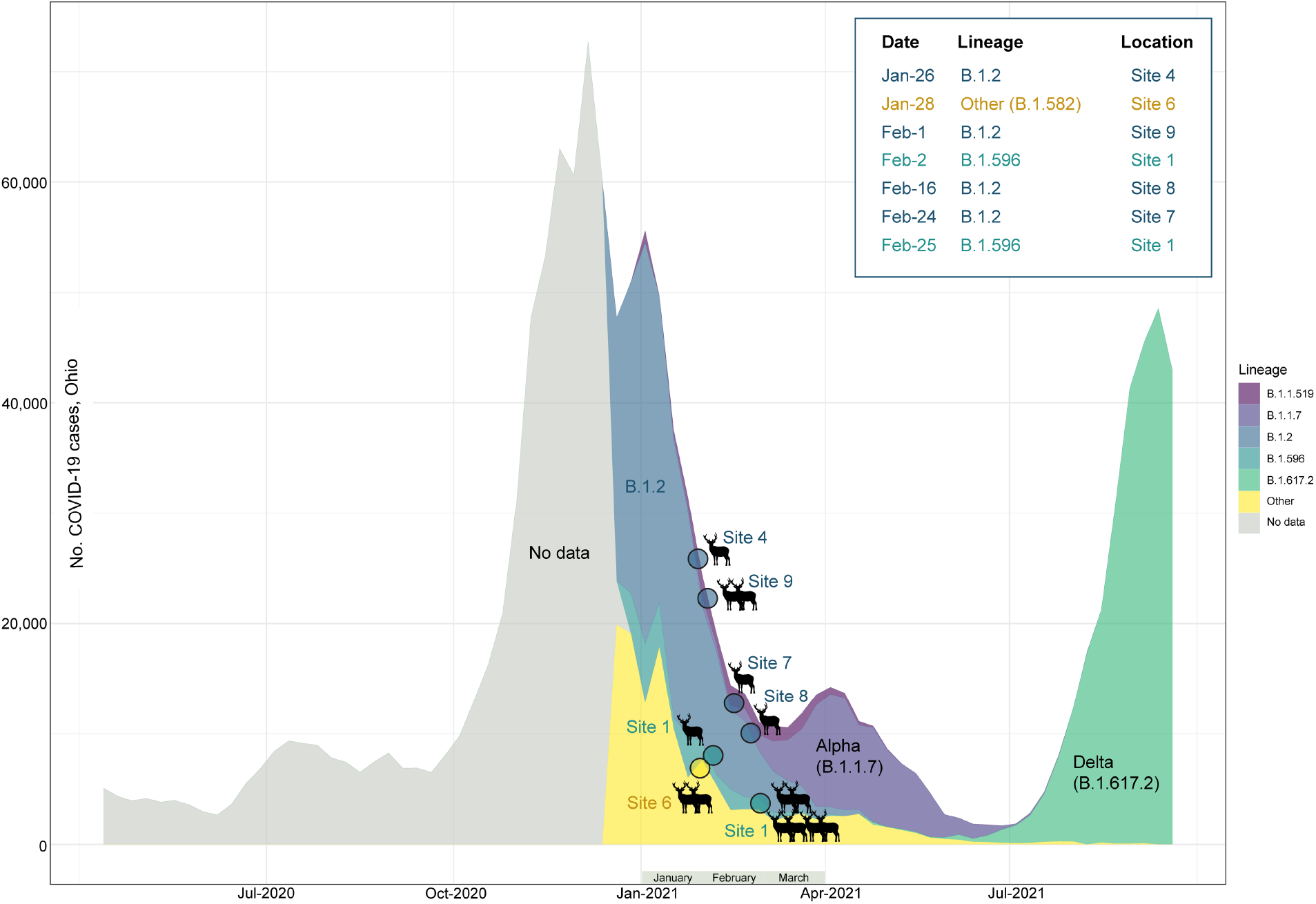
Timing of SARS-CoV-2 waves in Ohio, USA. The number of weekly COVID-19 cases in humans in Ohio is presented from April 2020 – September 2021 (y-axis). The proportion of viruses sequenced each week in Ohio for routine surveillance that are assigned to one of five Pango lineages (or “Other”) is indicated by shading. Genomic data prior to December 2020 was too sparse to reliably estimate lineage frequency. The shading and placement of each circle indicates the timing and lineage of SARS-CoV-2 viruses found in white-tailed deer at seven time points in this study.

Although B.1.2 was identified in deer at multiple sites, the phylogenetic analysis indicates that each site experienced a separate human-to-deer transmission event of a slightly genetically different B.1.2 virus (Figure 3). There was no evidence of B.1.2 virus transmitting across sites. Deer-to-deer transmission may have occurred within the three sites where multiple samples were sequenced: sites 1, 6, and 9 involving B.1.596, B.1.582, and B.1.2 lineages, respectively. It is impossible to know if deer-to-deer transmission occurred at the other 3 sites, as only one sample was successfully sequenced at each. The largest cluster was observed in site 1, where 7 deer sequences spanning two collection dates (02/02 and 02/25) fall together within clade B.1.596 (Figure 4). The 7 deer sequences in this clade contain 4 amino acid substitutions in ORF1 that were not observed in the most closely related human viruses despite having been observed in other viruses globally. Deletions were also observed in the ORF1 and spike proteins.

**Figure 3.**
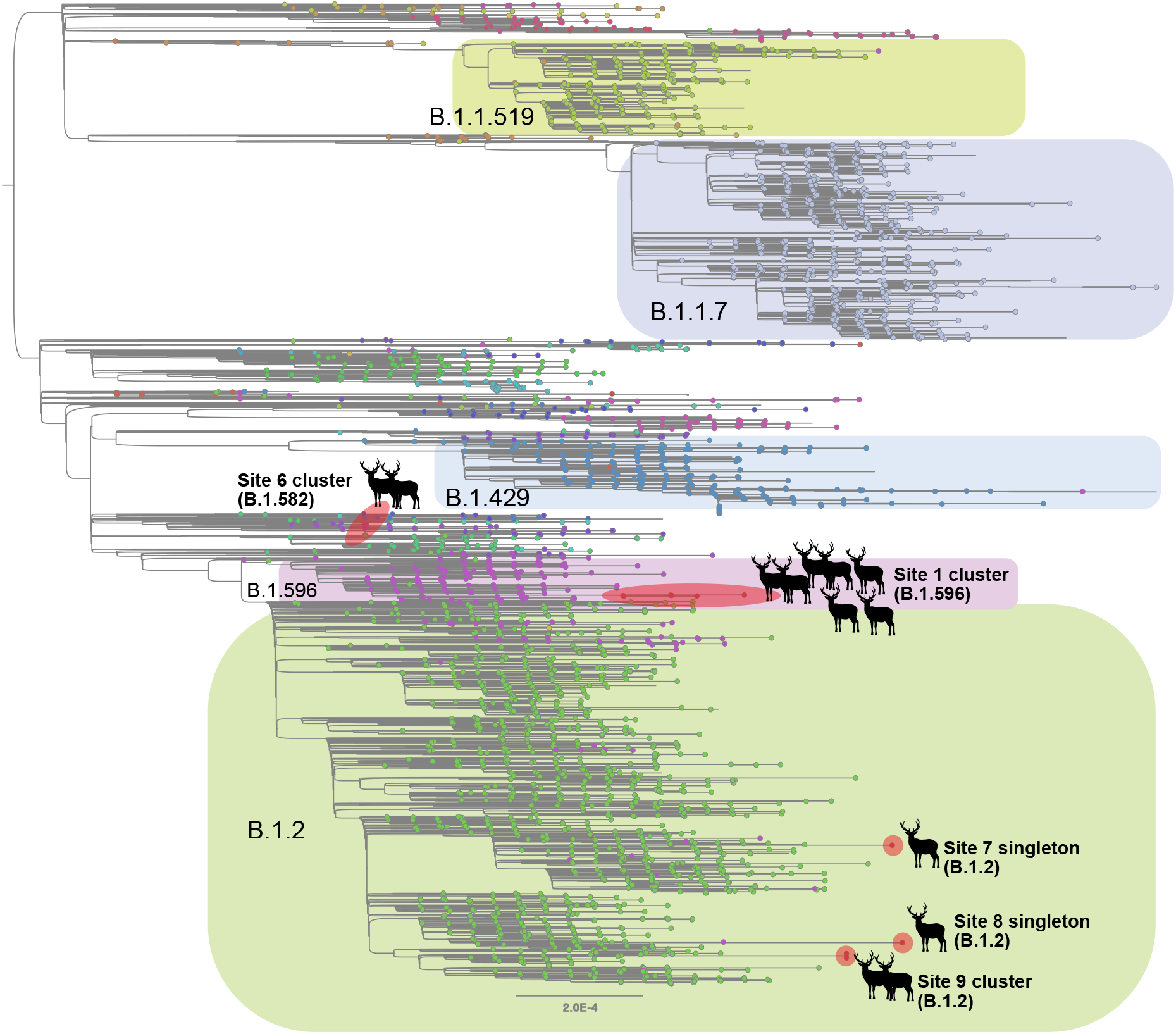
Evolutionary relationships of SARS-CoV-2 viruses in white-tailed deer. Maximum likelihood tree inferred for SARS-CoV-2 viruses in humans and white-tailed deer in Ohio during January – March 2021. Tips are shaded by Pango lineage and major lineages are labeled and boxed. Viruses found in white-tailed deer (clusters or singletons) are shaded red and labeled by location (site 4 not shown due to lower sequence coverage). All branch lengths drawn to scale.

**Figure 4.**
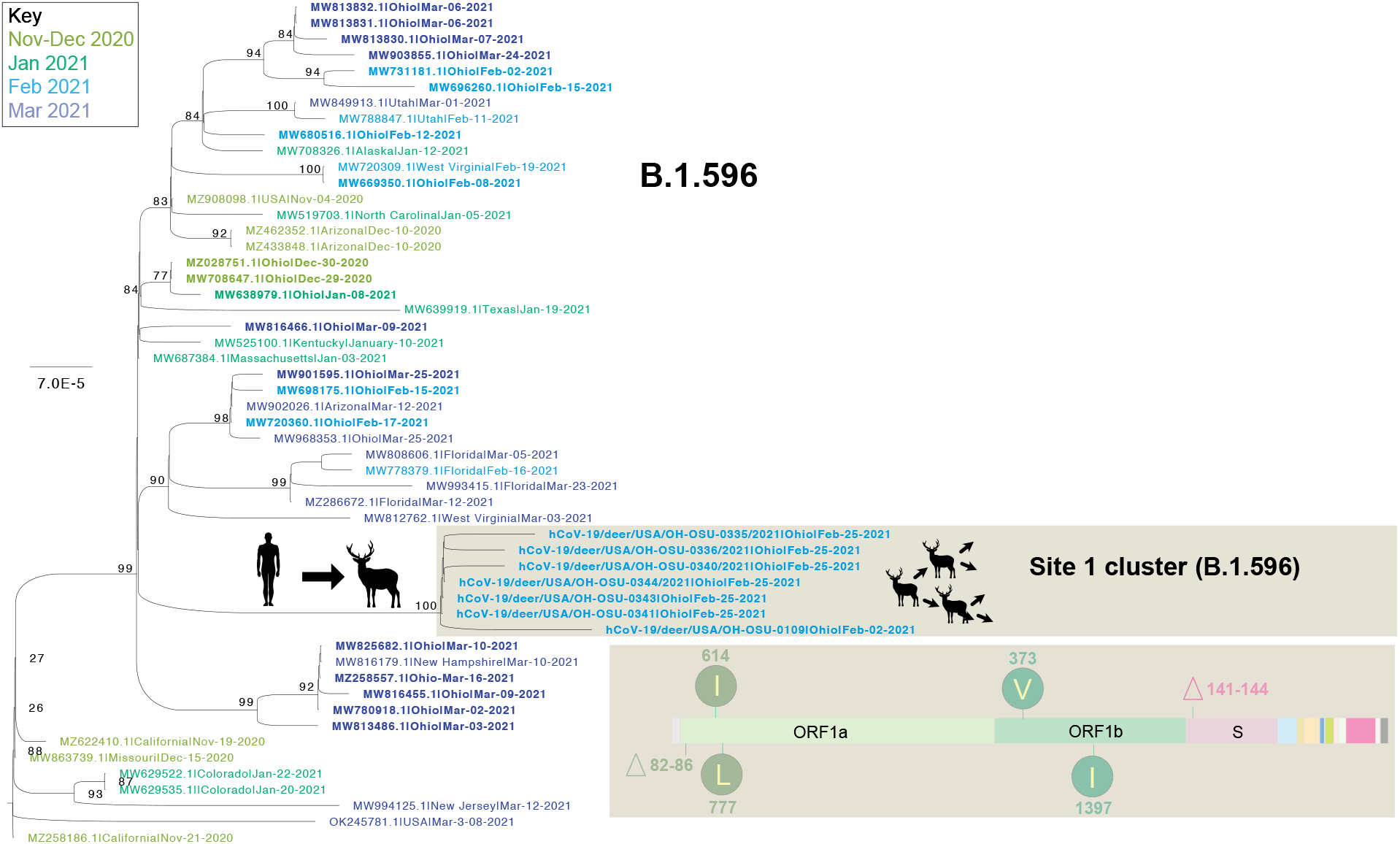
Evolutionary relationships of B.1.596 viruses in white-tailed deer. Maximum likelihood tree inferred for the cluster of 7 B.1.596 viruses found in white-tailed deer at site 1 and the 46 most closely related human B.1.596 viruses, all of which were observed in the United States during November 2020 – March 2021. Human viruses are shaded by month of collection. Cartoons indicate the branch where human-to-deer transmission may have occurred, followed by likely deer-to-deer transmission within site 1. Amino acid changes observed in the 7 B.1.596 deer viruses are listed. Bootstrap values are provided for key nodes and all branch lengths are drawn to scale.

## Discussion

Free-ranging white-tailed deer are distributed broadly across urban, suburban, and rural environments in the United States, and can live at densities of greater than 45 deer per square mile in some areas (*22*). Ohio is home to >600,000 free-ranging white-tailed deer (*23*) and another 440 commercial deer farms (*24*). Direct or indirect human-deer interaction occurs in parks and backyards, on livestock and cervid farms, in wildlife hospitals, and through hunting. The prevalence, distribution, and genetic variation of SARS-CoV-2 strains identified in free-ranging deer across Ohio raises a series of questions. How is the virus directly or indirectly transmitted from humans to deer? How extensively and efficiently are free-ranging deer transmitting the virus within or between species, in Ohio or other US states? Could viral adaptions occur in deer as has been observed in farmed mink? Could white-tailed deer serve as a reservoir for future spillovers of SARS-CoV-2 to humans or to other susceptible species?

The results from this study indicate that there have been multiple independent introductions of SARS-CoV-2 into deer occurring at different times and different locations around northeast Ohio. How these deer acquired SARS-CoV-2 from humans remains unclear. Our results are consistent with at least six human-to-deer transmission events occurring during Ohio’s major pandemic wave last winter, which peaked from November 2020 – January 2021. Although it is difficult to date precisely when these human-to-deer transmissions occurred, human-to-deer transmissions may be most likely during the weeks of highest virus circulation in humans. A similar temporal association was observed previously during the 2009 H1N1 influenza pandemic when the vast majority of transmission events from humans to swine and other species occurred during times of peak pandemic activity in humans. While high quality sequence coverage across the entire genome proved challenging for these post-mortem samples, it is likely that the presence of inhibitors within samples and/or sample degradation between collection and testing may have occurred. A comprehensive surveillance program in the future could provide additional data to help resolve remaining questions. Additionally, the recent wave of Delta variant infections in the human population likely represents another opportunity for SARS-CoV-2 to spillover into white-tailed deer. Sequencing of samples from the fall of 2021 will also determine whether the lineages seeded in deer last winter have continued to circulate.

It may be unsurprising that deer in urban sites were at higher risk for infection in our study. While it remains unknown how human-to-deer transmission occurs mechanistically, urban settings provide greater opportunities for direct and indirect contact with human-contaminated sources (e.g. trash, backyard feeders, bait stations), as well as contact with waterways that could be contaminated by multiple sources (*11, 25*). Viable SARS-CoV-2 is shed in human stool and has been demonstrated in wastewater (*26, 27*). Viable SARS-CoV-2 can be isolated from tap water and wastewater under experimental conditions (*26*). SARS-CoV-2 RNA has been identified in urban runoff in Ohio, which is useful for early detection systems. Virus isolation was not attempted and it is unknown whether SARS-CoV-2 was viable in these samples and capable of transmitting to other species (*28*). The recent detection of genetically disticnt SARS-CoV-2 virus fragments in New York City wastewater introduces the possibility that SARS-CoV-2 is transmitting cryptically in rodents (*29*). Contaminated waterways were also suggested as potential sources of infection for the free-ranging mink that tested positive for SARS-CoV-2 in Spain (*19*). Another question is how the virus transmits mechanistically between deer, whether by aerosols, direct contact, or environmentally. Prospective sequencing of samples from these deer populations will help determine whether the virus transmits long-term or only transiently in deer.

As SARS-CoV-2 continues to evolve it will be important to determine whether particular virus lineages are more associated with infection, transmission, disease, and persistence in deer. Although free-ranging white-tailed deer are highly susceptible to SARS-CoV-2 infection, it is unknown whether this leads to clinical disease. Two previous experimental studies reported subclinical infections in white-tailed deer challenged with SARS-CoV-2 (*7, 16*). The increased rate of infection in males found in this study could reflect differential susceptibility between sexes, but could also be explained by sex-linked differences in behavior that influence disease transmission. Male white-tailed deer also have a higher prevalence of chronic wasting disease and tuberculosis, which has been attributed to larger male home ranges, increased movement and contact with other deer during breeding season (fall/winter), and dynamic male social group composition and size (*30*). Monitoring SARS-CoV-2 evolution in wildlife populations, including white-tailed deer, mustelids, rodents, and other susceptible species will be critical to future COVID-19 management and mitigation efforts.

## References and Notes

References (*31–33*) in Supplementary Materials.

## Acknowledgments

Hannah Cochran, Elizabeth Ohl, Amber Cleggett, Sydney Treglia, Amanda M. Williams, Francesca Savona, Jacob W. Smith, Dakota Sizemore, and the Cleveland Metroparks Deer Management Team

## Funding

The Ohio State University Infectious Diseases Institute, Spring 2020 Interdisciplinary Seed Grant Program. eSCOUT--Environmental Surveillance for COVID-19 in Ohio: Understanding Transmission (VLH, PMD, JMN, DL, SG, LS, ASB)

Centers of Excellence for Influenza Research and Surveillance, National Institute of Allergy and Infectious Diseases, National Institutes of Health, Department of Health and Human Services, under contract HHSN272201400006C (DSM, JMN, DH, ASB) and HHSN272201400008C (MIN).

NIAID Intramural Research Program (MIN)

## Disclaimer

The content does not necessarily reflect the views or policies of the U.S. Government or imply endorsement of any products.

## Author contributions

Conceptualization: VLH, PAD, JMN, DL, SG, LS, SAF, ASB

Methodology: VLH, PAD, JMN, SAF, MIN, ASB

Investigation: PAD, JG, CM, DH, ME, RMT, MLK, KL, SRA, MT, ASB

Visualization: MIN, DSM, ASB

Funding acquisition: VLH, JMN, JAW, SAF, ASB

Project administration: VLH, PAD, JMN, ASB

Supervision: VLH, PAD, JG, MT, ASB

Writing – original draft: VLH, MIN, DSM, MLK, SAF, ASB

Writing – review & editing: PAD, CM, DH, JG, JMN, DL, JAW, SG, LS, ME, RMT, KL, SRA, MT

## Data and materials availability

All data are available in the main text or the supplementary materials.

## Supplementary Materials

Materials and Methods

Tables S1 - S5

