## Supplemental Materials for "SARS-CoV-2 infection in free-ranging white-tailed deer (*Odocoileus virginianus*)"

### Materials and Methods

**Sample collection**. Between January-March 2021, 360 free-ranging white-tailed deer originating from 9 study sites in northeast Ohio (USA) were euthanized as part of a deer population management program. Harvest occurred at locations that were baited for up to two weeks prior to each culling session, and additional deer were harvested opportunistically when they were observed away from the bait on a culling session day. Each day of the program, harvested deer carcasses were transported to a central processing point where samples were collected. Sample collectors wore gloves and a facemask. A nasal swab was collected from each deer and placed into a tube with brain heart infusion broth (BHIB). After collection, samples were immediately chilled on ice packs then transferred into a − 80°C freezer within 12 h where they remained until testing was initiated.

**Diagnostic testing**. Samples were initially tested using the Charité/Berlin (WHO) assay (31). Viral RNA was extracted from 200µl of BHIB using Omega Bio-tek Mag-Bind Viral DNA/RNA kit (cat #M6246-03). Xeno Internal Control (Life Technologies cat # A29763) was included in the extraction to ensure the accuracy of negative results. Five microliters of extracted RNA was added to Path-ID MPX One-Step Kit master mix (Life Technologies cat# 4442135) containing 12.5µl 2x Multiplex RT-PCR buffer, 2.5µl enzyme mix, 1.5µl nuclease free water, 4.5µl E assay primer/probe panel (Integrated DNA Technologies, Inc. cat #1006804), and 1µl XENO VIC Internal Control Assay (Life Technologies cat# A29765) for each sample. The cycling parameters for the real-time, reverse-transcriptase PCR (rRT-PCR) were 48ºC for 10 min, 95ºC 10 min, 45 cycles of 95ºC 15 sec and 58ºC 45 sec. Samples with a cycle threshold (Ct) of $\leq$40 were considered positive. If the E assay was positive, the RdRp confirmatory and discriminatory assays were completed using the above master mix formulation and thermocycler parameters, replacing the E assay primer/ probe panel with the confirmatory and discriminatory primer/probe panel (Integrated DNA Technologies, Inc. cat #s 10006805 and 10006806). The RNA from all samples that tested positive by the E assay, was retested by the CDC rRT-PCR protocol (32) Samples that were 2019-nCoV N1 and N2 positive were classified as presumptive positive.

**Genomic sequencing**. Representative presumptive positive samples (n=76) were sent to the National Veterinary Services Laboratories (NVSL) for confirmatory rRT-PCR testing using the CDC protocol and whole genome sequencing. Viral RNA was amplified by PCR (protocol available upon request) and cDNA libraries were prepared using the Nextera XT DNA Sample Preparation Kit according to manufacturer instructions. Sequencing was performed using the 500 cycle MiSeq Reagent Kit v2. Sequences were assembled using IRMA v0.6.7 and DNAStar SeqMan NGen v14.0.1. Additional sequencing was attempted at Ohio State's Applied Microbiology Services Laboratory using a modified ARTIC V3 method (33). Extracted RNA was reverse transcribed and amplified by polymerase chain reaction (PCR) with the ARTIC SARS-CoV-2 FS Library Prep Kit (New England Biolabs, Ipswich MA) per manufacturer's recommended protocol. Amplified products were converted into Illumina sequencing libraries using the RNA Prep with Enrichment (L) Tagmentation Kit protocol (Illumina, San Diego, CA, USA) with unique dual indexes and 10 cycles of TagPCR. Sequencing libraries were pooled and quantified using ProNex NGS Library Quant Kit (NG1201, Promega Co. Madison, WI). 650pM libraries were loaded on P2 sequencing cartridges and analyzed with the NextSeq2000 (lllumina) with 2x101bp cycles. Data were transmitted to the BaseSpace Cloud platform (Illumina) and converted to FASTQ file format using DRAGEN FASTQ Generation v3.8.4 (Illumina).

DRAGEN COVID Lineage app v3.5.3 (Illumina) was used to align sequence data and produce quality metrics and consensus genome sequences. Whole-genome sequences are available on GISAID for all 14 deer viruses sequenced for this study (Table S5).

**Data analysis**. Pangolin v3.1.11, 2021-09-17 was used to assign lineage. Prevalence was estimated using the number of presumptive positive nasal swabs based upon the final CDC rRT-PCR results. Prevalence estimates, confidence intervals, and other descriptive statistics were calculated using STATA 14.2 (StataCorp LLC).

**Phylogenetic analysis**. First, a background dataset was compiled from GISAID that included all SARS-CoV-2 sequences available from humans in Ohio, USA during the study period (January 1 – March 31, 2021), downloaded on September 27, 2021 (n = 4,801 sequences). To our knowledge, these are the first SARS-CoV-2 viruses sequenced from white-tailed deer globally and no additional sequences from white-tailed deer were available in any public repository for comparison. Pangolin was used to assign a lineage to each human virus. In total, 102 lineages were identified in this data set, with the most common being B.1.2 (n = 1766), B.1.1.7 (n = 833), 411 (n = B.1.1.519), B.1.429 (n = 307), and B.1.596 (n = 274). The dataset was aligned using NextClade with Wuhan-Hu-1 as a reference. The alignment was manually trimmed at the 5' and 3' ends. The final alignment included only coding regions and was manually edited to be in frame, with stop codons present only at the terminus of genes. A phylogenetic tree was inferred from this data set using maximum likelihood methods available in IQ-TREE version v1.6.12 with a GTR+G model of nucleotide substitution and 1,000 bootstrap replicates, using the high-performance computational capabilities of the Biowulf Linux cluster at the National Institutes of Health (http://biowulf.nih.gov). The inferred tree was visualized in FigTree v1.4.4. Outlier sequences were removed with long branch lengths and incongruence between genetic divergence and sampling date, as assessed using TempEst, typically arising from poor sequence coverage. One of the 14 sequences obtained from deer in our study (hCoV-19/deer/USA/OH-OSU-0025/2021, site 4) was lower in coverage and had a very long branch length and was excluded from the final phylogenetic analysis. To examine the evolutionary origins of the cluster of 7 B.1.596 viruses obtained from deer at site 1 in greater detail, a second phylogenetic tree was inferred that included all B.1.596 sequences available globally from NCBI's GenBank (n = 5,586), nearly all (99.8%) from the United States, using similar methods as above. The NVSL vSNP pipeline ((https://giithub.com/USDA-VS/vSNP) was applied for SNP based phylogenetic analysis using Wuhan-Hu-1 (NC_045512) as a reference.

**Epidemiological data**. The epidemiological curve of SARS-CoV-2 cases in Ohio from April 2020 to September 2020 was generated using the number of daily reported COVID-19 cases in the state of Ohio (all age groups), available from the US Centers for Disease Control and Prevention (https://data.cdc.gov/Case-Surveillance/COVID-19-Case-Surveillance-Public-Use-Data-with-Ge/n8mc-b4w4). All SARS-CoV-2 genetic sequences from Ohio were downloaded from GISAID on October 8, 2021 (n = 18,052) to estimate the proportion of viruses belonging to different Pango lineages during each week of the epidemic. To account for the intensity of surveillance not being even over time the number of viruses per lineage per week was normalized against the epidemiological curve derived from COVID-19 case counts and visualized using XX in R. To further minimize biases only sequences categorized in the GISAID submission as obtained using a "baseline surveillance" sampling strategy were included in the analysis. The dataset was further trimmed to include only submissions with complete collection dates and sufficient coverage to assign a Pango lineage, resulting in a final dataset of 9,947 sequences from Ohio. For simplicity sub-lineages of B.1.617.2 (e.g., AY.3) were consolidated into the Delta category and sub-lineages of B.1.1.7 (e.g., Q.3) were consolidated into the Alpha category. Baseline surveillance data prior to December 20, 2020 was too thinly sampled to reliably estimate the proportion of viruses from different lineages and was shaded grey to indicate insufficient data.

**Table S1. rRT-PCR testing results.** The cycle threshold (Ct) value results for the E assay screen, N1, and N2 rRT-PCR targets are shown for the 360 nasal swabs collected from white-tailed deer as a part of this study. Final SARS-CoV-2 rRT-PCR result was considered presumptive positive if Ct values for all three targets were less ≤ 40. For the 76 samples sent to NVSL, confirmatory rRT-PCR results are listed in addition to the samples for which whole genome sequencing (WGS) was successfully completed. See Table S5 for GISAID accession numbers for the 14 sequences generated.

| Sample ID | E assay | N1 assay | N2 assay | Presumptive PCR results | NVSL PCR results | WGS Completed |
|---|---|---|---|---|---|---|
| 1 | 30.20 | 30.72 | 29.13 | Positive | Positive | |
| 2 | 28.50 | 29.18 | 27.65 | Positive | Positive | |
| 3 | 29.33 | 29.65 | 28.39 | Positive | Positive | |
| 4 | 32.18 | 31.14 | 30.33 | Positive | | |
| 5 | Negative | | | Negative | | |
| 6 | Negative | | | Negative | | |
| 7 | 28.38 | 29.47 | 28.09 | Positive | Positive | |
| 8 | 28.63 | 29.70 | 27.42 | Positive | Positive | |
| 9 | 28.14 | 28.68 | 26.85 | Positive | Positive | |
| 10 | 17.56 | Negative | Negative | Negative | | |
| 11 | 35.98 | 35.00 | 34.10 | Positive | | |
| 12 | Negative | | | Negative | | |
| 13 | Negative | | | Negative | | |
| 14 | Negative | | | Negative | | |
| 15 | Negative | | | Negative | | |
| 16 | Negative | | | Negative | | |
| 17 | Negative | | | Negative | | |
| 18 | Negative | | | Negative | | |
| 19 | Negative | | | Negative | | |
| 20 | Negative | | | Negative | | |
| 21 | Negative | | | Negative | | |
| 22 | Negative | | | Negative | | |
| 23 | Negative | | | Negative | | |
| 24 | Negative | | | Negative | | |
| 25 | 23.67 | 24.76 | 22.73 | Positive | Positive | Yes |
| 26 | 31.06 | 29.41 | 31.27 | Positive | Positive | |
| 27 | Negative | | | Negative | | |
| 28 | Negative | | | Negative | | |
| 29 | Negative | | | Negative | | |
| 30 | Negative | | | Negative | | |
| 31 | 32.07 | 30.85 | 32.71 | Positive | | |
| 32 | Negative | | | Negative | | |
| 33 | 30.88 | 30.08 | 31.48 | Positive | | |

| | | | | | | |
|---|---|---|---|---|---|---|
| 34 | 30.57 | 29.63 | 31.76 | Positive | Positive | |
| 35 | Negative | | | Negative | | |
| 36 | Negative | | | Negative | | |
| 37 | Negative | | | Negative | | |
| 38 | Negative | | | Negative | | |
| 39 | Negative | | | Negative | | |
| 40 | Negative | | | Negative | | |
| 41 | Negative | | | Negative | | |
| 42 | Negative | | | Negative | | |
| 43 | Negative | | | Negative | | |
| 44 | Negative | | | Negative | | |
| 45 | Negative | | | Negative | | |
| 46 | Negative | | | Negative | | |
| 47 | Negative | | | Negative | | |
| 48 | Negative | | | Negative | | |
| 49 | Negative | | | Negative | | |
| 50 | 28.77 | 29.91 | 33.86 | Positive | Positive | |
| 51 | Negative | | | Negative | | |
| 52 | Negative | | | Negative | | |
| 53 | Negative | | | Negative | | |
| 54 | Negative | | | Negative | | |
| 55 | Negative | | | Negative | | |
| 56 | Negative | | | Negative | | |
| 57 | 25.21 | 26.49 | 29.85 | Positive | Positive | Yes |
| 58 | 25.23 | 26.91 | 29.81 | Positive | Positive | Yes |
| 59 | Negative | | | Negative | | |
| 60 | Negative | | | Negative | | |
| 61 | Negative | | | Negative | | |
| 62 | Negative | | | Negative | | |
| 63 | Negative | | | Negative | | |
| 64 | Negative | | | Negative | | |
| 65 | Negative | | | Negative | | |
| 66 | Negative | | | Negative | | |
| 67 | Negative | | | Negative | | |
| 68 | 31.00 | 28.93 | 30.36 | Positive | Positive | |
| 69 | 30.49 | 28.82 | 30.42 | Positive | Positive | |
| 70 | 30.27 | 28.30 | 29.81 | Positive | Positive | |
| 71 | 31.64 | 30.08 | 32.68 | Positive | | |
| 72 | Negative | | | Negative | | |
| 73 | 28.84 | 27.08 | 28.49 | Positive | Positive | |
| 74 | 29.29 | 28.18 | 30.80 | Positive | Positive | |

| | | | | | | |
|---|---|---|---|---|---|---|
| 75 | 32.38 | 30.57 | 32.09 | Positive | | |
| 76 | Negative | | | Negative | | |
| 77 | Negative | | | Negative | | |
| 78 | 33.32 | 31.91 | 34.14 | Positive | | Yes |
| 79 | 24.55 | 25.10 | 26.74 | Positive | Positive | Yes |
| 80 | Negative | | | Negative | | |
| 81 | Negative | | | Negative | | |
| 82 | Negative | | | Negative | | |
| 83 | 32.97 | 30.96 | 33.44 | Positive | | |
| 84 | 31.61 | 29.91 | 31.04 | Positive | Positive | |
| 85 | Negative | | | Negative | | |
| 86 | Negative | | | Negative | | |
| 87 | Negative | | | Negative | | |
| 88 | 24.28 | 22.62 | 23.96 | Positive | Positive | |
| 89 | Negative | | | Negative | | |
| 90 | Negative | | | Negative | | |
| 91 | Negative | | | Negative | | |
| 92 | Negative | | | Negative | | |
| 93 | Negative | | | Negative | | |
| 94 | Negative | | | Negative | | |
| 95 | Negative | | | Negative | | |
| 96 | Negative | | | Negative | | |
| 97 | Negative | | | Negative | | |
| 98 | Negative | | | Negative | | |
| 99 | Negative | | | Negative | | |
| 100 | Negative | | | Negative | | |
| 101 | Negative | | | Negative | | |
| 102 | Negative | | | Negative | | |
| 103 | Negative | | | Negative | | |
| 104 | Negative | | | Negative | | |
| 105 | Negative | | | Negative | | |
| 106 | Negative | | | Negative | | |
| 107 | Negative | | | Negative | | |
| 108 | Negative | | | Negative | | |
| 109 | 18.57 | 16.79 | 18.75 | Positive | Positive | Yes |
| 110 | 39.87 | Negative | Negative | Negative | | |
| 111 | Negative | | | Negative | | |
| 112 | Negative | | | Negative | | |
| 113 | Negative | | | Negative | | |
| 114 | Negative | | | Negative | | |
| 115 | Negative | | | Negative | | |

| | | | | | | |
|---|---|---|---|---|---|---|
| 116 | 33.17 | 34.38 | 35.19 | Positive | | |
| 116 | Negative | | | Negative | | |
| 117 | Negative | | | Negative | | |
| 118 | Negative | | | Negative | | |
| 119 | Negative | | | Negative | | |
| 120 | Negative | | | Negative | | |
| 121 | Negative | | | Negative | | |
| 122 | Negative | | | Negative | | |
| 123 | Negative | | | Negative | | |
| 124 | Negative | | | Negative | | |
| 125 | Negative | | | Negative | | |
| 126 | Negative | | | Negative | | |
| 127 | Negative | | | Negative | | |
| 128 | Negative | | | Negative | | |
| 129 | Negative | | | Negative | | |
| 131 | 32.96 | 32.89 | 35.00 | Positive | | |
| 132 | 29.74 | 29.62 | 31.97 | Positive | Positive | |
| 133 | Negative | | | Negative | | |
| 134 | Negative | | | Negative | | |
| 135 | 27.64 | 27.23 | 29.24 | Positive | Positive | |
| 136 | 29.25 | 28.67 | 30.53 | Positive | Positive | |
| 137 | 27.08 | 26.62 | 29.62 | Positive | Positive | |
| 138 | Negative | | | Negative | | |
| 139 | Negative | | | Negative | | |
| 140 | Negative | | | Negative | | |
| 141 | 26.17 | 26.74 | 28.55 | Positive | Positive | Yes |
| 142 | 27.91 | 31.20 | 32.77 | Positive | | |
| 143 | 31.25 | 27.44 | 29.30 | Positive | Positive | |
| 144 | Negative | | | Negative | | |
| 145 | Negative | | | Negative | | |
| 146 | Negative | | | Negative | | |
| 147 | Negative | | | Negative | | |
| 148 | Negative | | | Negative | | |
| 149 | Negative | | | Negative | | |
| 150 | Negative | | | Negative | | |
| 151 | Negative | | | Negative | | |
| 152 | Negative | | | Negative | | |
| 153 | Negative | | | Negative | | |
| 154 | 30.59 | 29.24 | 30.81 | Positive | Positive | |
| 155 | Negative | | | Negative | | |
| 156 | Negative | | | Negative | | |

| | | | | | |
|---|---|---|---|---|---|
| 157 | Negative | | | Negative | |
| 158 | 28.18 | 26.59 | 28.51 | Positive | Positive |
| 159 | Negative | | | Negative | |
| 160 | Negative | | | Negative | |
| 161 | Negative | | | Negative | |
| 162 | 31.73 | 30.44 | 32.82 | Positive | |
| 163 | Negative | | | Negative | |
| 164 | Negative | | | Negative | |
| 165 | 34.03 | 31.20 | 33.93 | Positive | |
| 166 | Negative | | | Negative | |
| 167 | Negative | | | Negative | |
| 168 | 31.72 | 29.53 | 32.28 | Positive | Positive |
| 169 | Negative | | | Negative | |
| 170 | Negative | | | Negative | |
| 171 | Negative | | | Negative | |
| 172 | Negative | | | Negative | |
| 173 | 31.21 | 29.55 | 34.14 | Positive | Positive |
| 174 | Negative | | | Negative | |
| 175 | Negative | | | Negative | |
| 176 | Negative | | | Negative | |
| 177 | Negative | | | Negative | |
| 178 | Negative | | | Negative | |
| 179 | 32.78 | 31.17 | 33.66 | Positive | |
| 180 | Negative | | | Negative | |
| 181 | Negative | | | Negative | |
| 182 | Negative | | | Negative | |
| 183 | 30.08 | 28.43 | 30.49 | Positive | Positive |
| 184 | Negative | | | Negative | |
| 185 | Negative | | | Negative | |
| 186 | Negative | | | Negative | |
| 187 | 31.86 | 32.98 | 31.25 | Positive | |
| 188 | 29.79 | 31.02 | 29.21 | Positive | Positive |
| 189 | 34.93 | 32.37 | 33.62 | Positive | |
| 190 | 26.63 | 28.80 | 25.94 | Positive | Positive |
| 191 | Negative | | | Negative | |
| 192 | Negative | | | Negative | |
| 193 | Negative | | | Negative | |
| 194 | 30.71 | 33.63 | 31.48 | Positive | |
| 195 | 31.71 | 31.93 | 30.36 | Positive | |
| 196 | Negative | | | Negative | |
| 197 | Negative | | | Negative | |

| | | | | | | |
|---|---|---|---|---|---|---|
| 198 | Negative | | | Negative | | |
| 199 | 27.05 | 27.23 | 24.82 | Positive | Positive | |
| 200 | Negative | | | Negative | | |
| 201 | Negative | | | Negative | | |
| 202 | Negative | | | Negative | | |
| 203 | Negative | | | Negative | | |
| 204 | Negative | | | Negative | | |
| 205 | 32.91 | 33.38 | 31.59 | Positive | | |
| 206 | 34.34 | 33.60 | 34.12 | Positive | | |
| 207 | 31.95 | 34.06 | 32.82 | Positive | | |
| 208 | 32.36 | 33.81 | 31.63 | Positive | | |
| 209 | Negative | | | Negative | | |
| 210 | 27.19 | 28.62 | 27.17 | Positive | Positive | |
| 211 | 29.37 | 31.00 | 29.87 | Positive | Positive | |
| 212 | 27.35 | 30.96 | 28.66 | Positive | Positive | Yes |
| 213 | Negative | | | Negative | | |
| 214 | 31.04 | 33.32 | 32.51 | Positive | | |
| 215 | Negative | | | Negative | | |
| 216 | 27.16 | 28.98 | 27.32 | Positive | Positive | |
| 289 | 24.41 | 25.61 | 23.65 | Positive | Positive | |
| 290 | 29.59 | 31.49 | 29.77 | Positive | Positive | |
| 291 | 27.27 | 29.00 | 27.61 | Positive | Positive | |
| 292 | 31.20 | 32.56 | 31.23 | Positive | | |
| 293 | 30.52 | 32.28 | 30.45 | Positive | | |
| 294 | Negative | | | Negative | | |
| 295 | Negative | | | Negative | | |
| 296 | Negative | | | Negative | | |
| 297 | 30.93 | 32.90 | 32.40 | Positive | | |
| 298 | 29.05 | 30.66 | 28.76 | Positive | Positive | |
| 299 | 30.53 | 31.15 | 29.63 | Positive | Positive | |
| 300 | 30.21 | 30.69 | 28.87 | Positive | Positive | |
| 301 | 33.98 | 34.07 | 33.82 | Positive | | |
| 302 | 26.93 | 28.71 | 26.79 | Positive | Positive | |
| 303 | 29.31 | 31.12 | 29.36 | Positive | Positive | |
| 304 | 31.78 | 32.38 | 31.33 | Positive | | |
| 305 | 32.16 | 32.32 | 31.25 | Positive | | |
| 306 | 33.13 | 33.62 | 31.63 | Positive | | |
| 307 | 29.08 | 30.35 | 28.84 | Positive | Positive | |
| 308 | Negative | | | Negative | | |
| 309 | 30.71 | 31.51 | 29.66 | Positive | Positive | |
| 310 | 30.21 | 31.12 | 31.19 | Positive | | |

| | | | | | | |
|---|---|---|---|---|---|---|
| 311 | 35.92 | 32.75 | 29.34 | Positive | Positive | |
| 312 | 32.33 | Negative | 35.51 | Negative | | |
| 313 | 33.80 | 34.91 | 35.00 | Positive | | |
| 314 | 28.69 | 29.52 | 27.96 | Positive | Positive | |
| 315 | Negative | | | Negative | | |
| 316 | 28.36 | 30.04 | 28.89 | Positive | Positive | |
| 317 | 29.36 | 30.56 | 29.19 | Positive | Positive | |
| 318 | 29.56 | 30.89 | 29.33 | Positive | Positive | |
| 319 | Negative | | | Negative | | |
| 320 | Negative | | | Negative | | |
| 321 | 29.58 | 31.05 | 30.02 | Positive | | |
| 322 | 26.91 | 28.24 | 26.78 | Positive | Positive | |
| 323 | 30.69 | 31.51 | 29.72 | Positive | Positive | |
| 324 | 36.35 | 35.40 | 33.01 | Positive | | |
| 325 | Negative | | | Negative | | |
| 326 | Negative | | | Negative | | |
| 327 | Negative | | | Negative | | |
| 328 | 31.14 | 33.30 | 31.82 | Positive | | |
| 329 | Negative | | | Negative | | |
| 330 | Negative | | | Negative | | |
| 331 | Negative | | | Negative | | |
| 332 | Negative | | | Negative | | |
| 333 | Negative | | | Negative | | |
| 334 | Negative | | | Negative | | |
| 335 | 26.18 | 27.70 | 25.88 | Positive | Positive | Yes |
| 336 | 14.01 | 12.88 | 12.71 | Positive | Positive | Yes |
| 337 | 32.89 | 35.32 | 23.66 | Positive | Positive | |
| 338 | Negative | | | Negative | | |
| 339 | Negative | | | Negative | | |
| 340 | 22.03 | 22.82 | 16.90 | Positive | Positive | Yes |
| 341 | 26.75 | 29.46 | 35.48 | Positive | Positive | Yes |
| 342 | 33.26 | 35.75 | 27.68 | Positive | Negative | |
| 343 | 16.58 | 16.45 | 35.39 | Positive | Positive | Yes |
| 344 | 26.81 | 30.40 | 27.75 | Positive | Positive | Yes |
| 345 | Negative | | | Negative | | |
| 346 | Negative | | | Negative | | |
| 347 | Negative | | | Negative | | |
| 348 | Negative | | | Negative | | |
| 349 | Negative | | | Negative | | |
| 350 | Negative | | | Negative | | |
| 351 | Negative | | | Negative | | |

| | | | | | |
|---|---|---|---|---|---|
| 352 | Negative | | | Negative | |
| 353 | Negative | | | Negative | |
| 354 | Negative | | | Negative | |
| 355 | Negative | | | Negative | |
| 356 | Negative | | | Negative | |
| 357 | Negative | | | Negative | |
| 358 | Negative | | | Negative | |
| 359 | Negative | | | Negative | |
| 360 | Negative | | | Negative | |
| 361 | Negative | | | Negative | |
| 362 | Negative | | | Negative | |
| 363 | Negative | | | Negative | |
| 364 | Negative | | | Negative | |
| 365 | Negative | | | Negative | |
| 366 | Negative | | | Negative | |
| 367 | Negative | | | Negative | |
| 368 | Negative | | | Negative | |
| 369 | Negative | | | Negative | |
| 370 | Negative | | | Negative | |
| 371 | Negative | | | Negative | |
| 372 | 30.97 | 29.60 | 28.14 | Positive | Positive |
| 373 | 34.04 | 33.32 | 31.67 | Positive | |
| 374 | 36.32 | 35.41 | 34.10 | Positive | |
| 375 | 33.41 | 31.98 | 30.86 | Positive | |
| 376 | 31.87 | 29.96 | 28.50 | Positive | Positive |
| 377 | Negative | | | Negative | |
| 378 | Negative | | | Negative | |
| 379 | 30.06 | 29.16 | 27.65 | Positive | Positive |
| 380 | Negative | | | Negative | |
| 381 | Negative | | | Negative | |
| 382 | Negative | | | Negative | |
| 383 | Negative | | | Negative | |
| 384 | Negative | | | Negative | |
| 385 | 30.27 | 28.54 | 27.23 | Positive | Positive |
| 386 | 32.96 | 31.57 | 30.07 | Positive | |
| 387 | Negative | | | Negative | |
| 388 | Negative | | | Negative | |
| 389 | Negative | | | Negative | |
| 390 | Negative | | | Negative | |
| 391 | Negative | | | Negative | |
| 392 | Negative | | | Negative | |

| | | | | | |
|---|---|---|---|---|---|
| 393 | 38.78 | Negative | Negative | Negative | |
| 394 | Negative | | | Negative | |
| 395 | Negative | | | Negative | |
| 396 | 30.38 | 28.80 | 27.69 | Positive | Positive |
| 397 | 33.92 | 31.20 | 30.46 | Positive | |
| 398 | 35.70 | 33.83 | 31.90 | Positive | |
| 399 | 33.86 | 31.30 | 29.84 | Positive | Positive |
| 400 | Negative | | | Negative | |
| 401 | 34.76 | 32.19 | 30.49 | Positive | |
| 402 | Negative | | | Negative | |
| 403 | Negative | | | Negative | |
| 404 | Negative | | | Negative | |
| 405 | 33.89 | 31.43 | 30.53 | Positive | |
| 406 | 33.75 | 31.70 | 30.15 | Positive | |
| 407 | 38.60 | 36.94 | 34.78 | Positive | |
| 408 | Negative | | | Negative | |
| 409 | 34.94 | 31.82 | 30.19 | Positive | |
| 410 | 32.70 | 30.26 | 28.42 | Positive | Positive |
| 411 | 31.82 | 29.50 | 28.25 | Positive | Positive |
| 412 | 35.97 | 35.09 | 37.68 | Positive | |
| 413 | 36.68 | 34.37 | 37.41 | Positive | |
| 414 | Negative | | | Negative | |
| 415 | Negative | | | Negative | |
| 416 | Negative | | | Negative | |
| 417 | Negative | | | Negative | |
| 418 | 36.91 | 37.89 | Negative | Negative | |
| 419 | 32.94 | 31.81 | 33.82 | Positive | Positive |
| 420 | Negative | | | Negative | |
| 421 | 34.66 | 33.37 | 38.09 | Positive | Positive |
| 422 | 31.70 | 31.91 | 33.18 | Positive | |
| 423 | 37.84 | 36.19 | Negative | Negative | |
| 424 | Negative | | | Negative | |
| 425 | 33.23 | 33.18 | 33.41 | Positive | |
| 426 | 32.98 | 32.70 | 33.93 | Positive | |
| 427 | 36.00 | 33.47 | 31.81 | Positive | |
| 428 | 33.27 | 30.75 | 29.22 | Positive | Positive |
| 429 | 34.36 | 31.54 | 29.77 | Positive | Positive |
| 430 | 37.63 | 35.88 | 35.29 | Positive | |
| 431 | Negative | | | Negative | |
| 432 | Negative | | | Negative | |

**Table S2. SARS-CoV-2 prevalence stratified by site and sampling date.** Sample collection dates for all nine sites are shown indicating the number of nasal swabs collected from white-tailed deer on each date and the number of those swabs that were screened positive for SARS-CoV-2 using the WHO and Centers for Disease Control and Prevention rRT-PCR protocols in series. Overall estimated prevalence is shown with 95% confidence interval estimates (Clopper-Pearson exact). The 7 collection dates from which genetic sequences were obtained are in bold text.

| | Date of collection | Samples | Positive | Estimated Prevalence | Lower 95% CI | Upper 95% CI |
|---|---|---|---|---|---|---|
| Site 1 | **2021-02-02** | 41 | 2 | 5% | 1% | 17% |
| | **2021-02-25** | 33 | 8 | 24% | 11% | 42% |
| Site 2 | 2021-1-27 | 4 | 3 | 75% | 19% | 99% |
| | 2021-03-03 | 16 | 11 | 69% | 41% | 89% |
| Site 3 | 2021-03-02 | 24 | 8 | 33% | 16% | 55% |
| Site 4 | **2021-01-26** | 14 | 2 | 14% | 2% | 43% |
| | 2021-02-17 | 16 | 4 | 25% | 7% | 52% |
| Site 5 | 2021-01-25 | 16 | 8 | 50% | 25% | 75% |
| | 2021-02-09 | 8 | 5 | 63% | 24% | 91% |
| | 2021-03-08 | 6 | 4 | 67% | 22% | 96% |
| Site 6 | **2021-01-28** | 33 | 3 | 9% | 2% | 24% |
| | 2021-03-04 | 15 | 7 | 47% | 21% | 73% |
| Site 7 | 2021-02-22 | 20 | 5 | 25% | 9% | 49% |
| | 2021-02-23 | 19 | 8 | 42% | 20% | 67% |
| | **2021-02-24** | 50 | 36 | 72% | 58% | 84% |
| Site 8 | **2021-02-16** | 13 | 3 | 23% | 5% | 54% |
| Site 9 | **2021-02-01** | 22 | 12 | 55% | 32% | 76% |
| | 2021-03-01 | 10 | 0 | 0% | 0% | 31%* |

**Table S3. Covariate data recorded for each deer sample.** rRT-PCR positive samples are shown by group for categorical variables, p-values shown from Pearson's Chi2. The mean of each continuous variable is show for rRT-PCR positive and negative deer respectively, p-values shows from Wilcoxon-Mann-Whitney rank-sum test. Age was not recorded for 3 deer which tested negative for SARS-CoV-2.

Categorical

| Covariate | Category | Samples | Positive (%) | P-Value |
|---|---|---|---|---|
| Sex | | | | <0.0005 |
| | Male | 149 | 76 (51) | |
| | Female | 211 | 53 (25) | |
| Cull method | | | | 0.395 |
| | Treestand | 119 | 39 (33) | |
| | Opportunistic | 241 | 90 (37) | |

Continuous

| Covariate | Samples | Negative mean | Positive mean | P-Value |
|---|---|---|---|---|
| Mass (Kg) | 360 | 52.76 | 56.73 | 0.0056 |
| Age (years) | 357 | 2.11 | 2.05 | 0.9717 |

**Table S4. Proportion of SARS-CoV-2 viruses identified in Ohio in humans during January 1 – February 28, 2021.** Only viruses categorized as collected for baseline surveillance are included.

| Rank | Lineage | Percentage | Number |
|---|---|---|---|
| 1 | B.1.2 | 56.3% | 462 |
| 2 | B.1.596 | 8.3% | 68 |
| 3 | B.1.1.519 | 7.1% | 58 |
| 4 | B.1.429 | 3.4% | 28 |
| 5 | B.1.1.7 | 3.0% | 25 |
| 6 | B.1.234 | 2.7% | 22 |
| 7 | P.2 | 2.4% | 20 |
| 8 | B.1.311 | 1.6% | 13 |
| 9 | B.1.427 | 1.5% | 12 |
| 10 | B.1.400 | 1.2% | 10 |
| 11 | B.1.582 | 1.2% | 10 |
| 12 | B.1.1 | 1.1% | 9 |
| 13 | B.1.243 | 1.1% | 9 |
| 14 | B.1 | 0.7% | 6 |
| 15 | B.1.568 | 0.7% | 6 |
| 16 | B.1.595 | 0.6% | 5 |
| 17 | B.1.1.337 | 0.5% | 4 |
| 18 | B.1.240 | 0.5% | 4 |
| 19 | B.1.448 | 0.5% | 4 |
| 20 | B.1.1.207 | 0.4% | 3 |
| 21 | B.1.1.434 | 0.4% | 3 |
| 22 | B.1.265 | 0.4% | 3 |
| 23 | B.1.361 | 0.4% | 3 |
| 24 | B.1.438.4 | 0.4% | 3 |
| 25 | B.1.577 | 0.4% | 3 |
| 26 | B.1.588 | 0.4% | 3 |
| 27 | B.1.1.265 | 0.2% | 2 |
| 28 | B.1.298 | 0.2% | 2 |
| 29 | B.1.517 | 0.2% | 2 |
| 30 | B.1.565 | 0.2% | 2 |
| 31 | Q.8 | 0.2% | 2 |
| 32 | B.1.1.291 | 0.1% | 1 |
| 33 | B.1.1.316 | 0.1% | 1 |
| 34 | B.1.1.348 | 0.1% | 1 |
| 35 | B.1.110.3 | 0.1% | 1 |
| 36 | B.1.139 | 0.1% | 1 |
| 37 | B.1.324 | 0.1% | 1 |
| 38 | B.1.346 | 0.1% | 1 |
| 39 | B.1.404 | 0.1% | 1 |
| 40 | B.1.409 | 0.1% | 1 |
| 41 | B.1.493 | 0.1% | 1 |
| 42 | B.1.503 | 0.1% | 1 |
| 43 | B.1.564 | 0.1% | 1 |
| 44 | C.23 | 0.1% | 1 |
| 45 | C.31 | 0.1% | 1 |

**Table S5. Whole genome sequence GISAID data.** Sequences generated from original sample nasal swabs collected from white-tailed deer as a part of this study are shown with GISAID accession numbers, collection date, and the submitting laboratory indicating where the sequencing was completed.

| Virus name | Accession ID | Collection date | Submitting laboratory |
|---|---|---|---|
| hCoV-19/deer/USA/OH-OSU-0025/2021 | EPI_ISL_4878314 | 2021-01-26 | The Ohio State University Applied Microbiology Services Laboratory |
| hCoV-19/deer/USA/OH-OSU-0057/2021 | EPI_ISL_4847029 | 2021-01-28 | USDA National Veterinary Services Laboratories |
| hCoV-19/deer/USA/OH-OSU-0058/2021 | EPI_ISL_4847030 | 2021-01-28 | USDA National Veterinary Services Laboratories |
| hCoV-19/deer/USA/OH-OSU-0078/2021 | EPI_ISL_4878315 | 2021-02-01 | The Ohio State University Applied Microbiology Services Laboratory |
| hCoV-19/deer/USA/OH-OSU-0079/2021 | EPI_ISL_4847031 | 2021-02-01 | The Ohio State University Applied Microbiology Services Laboratory |
| hCoV-19/deer/USA/OH-OSU-0109/2021 | EPI_ISL_4878316 | 2021-02-02 | USDA National Veterinary Services Laboratories |
| hCoV-19/deer/USA/OH-OSU-0141/2021 | EPI_ISL_4847032 | 2021-02-16 | USDA National Veterinary Services Laboratories |
| hCoV-19/deer/USA/OH-OSU-0212/2021 | EPI_ISL_4847033 | 2021-02-24 | USDA National Veterinary Services Laboratories |
| hCoV-19/deer/USA/OH-OSU-0335/2021 | EPI_ISL_4878317 | 2021-02-25 | USDA National Veterinary Services Laboratories |
| hCoV-19/deer/USA/OH-OSU-0336/2021 | EPI_ISL_4878318 | 2021-02-25 | USDA National Veterinary Services Laboratories |
| hCoV-19/deer/USA/OH-OSU-0340/2021 | EPI_ISL_4878319 | 2021-02-25 | USDA National Veterinary Services Laboratories |
| hCoV-19/deer/USA/OH-OSU-0341/2021 | EPI_ISL_4878320 | 2021-02-25 | USDA National Veterinary Services Laboratories |
| hCoV-19/deer/USA/OH-OSU-0343/2021 | EPI_ISL_4878321 | 2021-02-25 | USDA National Veterinary Services Laboratories |
| hCoV-19/deer/USA/OH-OSU-0344/2021 | EPI_ISL_4878322 | 2021-02-25 | The Ohio State University Applied Microbiology Services Laboratory |
